# Deep Neurite Analysis Tool (DeNAT): A machine-learning framework for precise automated neurite outgrowth measurement

**DOI:** 10.1101/2025.09.18.676567

**Authors:** Manojkumar Kumaran, PS Athul Narayan, Yogesh Sahu, Soupayan Banerjee, Anisha S Menon, Shringika Soni, Ishwariya Venkatesh

**Author notes:** **Corresponding Author:** Ishwariya Venkatesh.

## Abstract

Accurate quantification of neurite sprouting after injury is a critical step in axon regeneration research. Yet it remains a major bottleneck, as the current gold standard relies on manual counting by multiple blinded observers. This process is slow, labor-intensive, and prone to variability. While some software can measure total neurite length, they aren’t made to specifically measure new growth in complicated images from real-life injury models, like the thoracic crush and pyramidotomy model. Existing software can measure total neurite length in culture, but it is not designed to capture new growth in complex images from injury models, such as thoracic crush or pyramidotomy. Crucially, these tools lack the ability to selectively analyze growth within user-defined regions, a key requirement for injury paradigms. To address this need, we developed the Deep Neurite Analysis Tool (DeNAT), an accessible deep-learning–based platform that automatically measures neurite outgrowth after injury. DeNAT allows users to define regions of interest to specifically quantify sprouting in images from common spinal cord injury paradigms. We benchmarked its performance against manual scoring and conventional automated approaches. DeNAT achieved 87 percent accuracy in detecting neurite sprouts relative to manual counts, while reducing variability and labor. By combining user-guided region selection with automated deep learning analysis, DeNAT offers an accurate, reproducible, and efficient solution for measuring neurite outgrowth in injury models.

## Introduction

The successful regeneration of axons after injury is central to restoring neuronal circuit function, yet it remains one of the greatest challenges in neuroscience. Unlike developmental growth, regenerating axons must navigate inhibitory environments and complex injury sites, making their measurement particularly difficult. In models such as pyramidotomy, where collateral sprouting of corticospinal axons is examined, accurate quantification of new growth is critical but inherently tricky due to overlapping fibers, dense tissue architecture, and region-specific differences in regeneration (Starkey et al., 2005). Traditionally, neurite outgrowth assays *in vitro* have provided a framework to study factors that promote or inhibit growth (Sakurai et al., 1997; Kleene et al., 2001; De Jaco et al., 2002; Fournier et al., 2002; Ahmed et al., 2006; Y. Zhang et al., 2007; Pool et al., 2008) with average neurite length per cell serving as the most common metric. However, when applied to regeneration models, particularly *in vivo*, quantification is still performed largely by manual counting, a method that is time-consuming, error-prone, and prone to variability across observers. Several automated methods for measuring neurite outgrowth have been developed, but they are almost exclusively designed for cultured neurons rather than regenerating axons *in vivo*. Commercial systems such as the IN Cell Analyzer (GE Healthcare) or ImageXpress (Molecular Devices) provide high-throughput analysis but are prohibitively expensive for many laboratories. Published tracing algorithms (Al-Kofahi et al., 2006; W. Zhang et al., 2007) can achieve detailed reconstructions, yet they require substantial computational expertise, limiting accessibility for most biologists. As a result, many groups still rely on manual or semi-manual neurite tracing (Meijering et al., 2004), a method that is slow, variable, and error-prone. Importantly, the widely used metrics from culture systems, such as average neurite length per cell (Sakurai et al., 1997; Kleene et al., 2001; De Jaco et al., 2002; Fournier et al., 2002; Ahmed et al., 2006; W. Zhang et al., 2007; Pool et al., 2008), do not translate well to complex injury paradigms like the pyramidotomy model, where regenerating corticospinal fibers must be quantified in dense tissue with region-specific sprouting. Despite pyramidotomy being one of the most common spinal cord injury models (Carmel et al., 2014; Chang et al., 2022; Chen et al., 2024; Fink and Cafferty, 2016; Ito et al., 2018, 2018; Kathe et al., 2014; Kauer et al., 2022; Niemeyer et al., 2000) there is currently no software explicitly designed to automate the analysis of regenerating fibers in this context.

To overcome the limitations of existing methods, we developed the Deep Neurite Analysis Tool (DeNAT), a freely available program that applies machine learning to automatically trace regenerating fibers in the pyramidotomy model. DeNAT is tailored to detect biologically meaningful sprouting within user-defined regions, making it well-suited for analyzing corticospinal tract plasticity. We compared its performance with two commonly used automated tools (Simple Neurite Tracer and NeuriteTracer) and with manual scoring using Zen software. DeNAT not only reproduced manual accuracy but also reduced analysis time from 45–60 minutes per image to just 1–2 minutes, a greater than 95 percent reduction in effort. By uniting biological relevance with speed and reproducibility, DeNAT offers researchers a practical and reliable solution for quantifying axon regeneration in the pyramidotomy paradigm.

## Methods

### Animals

All animal procedures were performed in accordance with the guidelines with prior approval from the Institutional Animal Ethics Committee (IAEC) at CSIR-CCMB. Wild-type C57BL/6J mice (both male and female, 8–12 weeks old) were used for all experiments. Animals were housed in a controlled environment with a 12-hour light/dark cycle and had free access to standard food and water.

### Surgical Procedures

All surgeries were conducted under aseptic conditions. Prior to each procedure, mice were anesthetized with a mixture of ketamine and xylazine [e.g., 100 mg/kg ketamine and 10 mg/kg xylazine, administered intraperitoneally (IP)]. The adequacy of anesthesia was confirmed by the absence of a pedal withdrawal reflex. After surgery, mice were administered an analgesic [e.g., Meloxicam at 5 mg/kg] and allowed to recover on a heating pad.

### Cortical Viral Injections

Adult C57BL/6J mice of both sexes (>12 weeks old, 20-25g) were used. Each mouse was placed in a stereotaxic apparatus. After exposing the skull, small holes were drilled over the sensorimotor cortex at two sites relative to Bregma on one side of the brain (in mm): (AP: 0, ML: +2), (AP: +1, ML: +2), (AP: 0, ML: -2), and (AP: +1, ML: -2). A Hamilton syringe was used to inject 0.6 µL of virus at a depth of 0.50–0.60 mm from the dural surface at a rate of [e.g., 0.05 µL/min]. After each injection, the syringe was left in place for 5 minutes to prevent backflow before being slowly withdrawn. The skin incision was closed with sutures.

### Pyramidotomy

Pyramidotomy was performed 7 days post-injection (Starkey et al., 2005). Briefly, following a ventral midline incision in the neck, the occipital bone was exposed. After puncturing the dura, the right pyramidal tract was transected using a very fine surgical blade. Animals were sacrificed 8 weeks after surgery. The spinal cord was dissected from the upper cervical region (C1–C4), fixed overnight in 4% paraformaldehyde (PFA) at 4°C, and washed in PBS. The tissue was then embedded in gelatin, re-fixed briefly, and cut into 50 µm transverse sections on a vibratome.

### Image Processing and Quantification

For the pyramidotomy model, three 50 μm transverse spinal cord sections per animal were selected for analysis, sampled from spinal levels C2 to C6 with 1.5 mm spacing. Image stacks were acquired on a Leica STED microscope using a 63× objective for pyramidotomy samples and were captured as tile scans (520 × 520 pixels) with 1 μm z-steps, a scan speed of 400 Hz, 40% laser power, and a detector gain of 1,250 V. For normalization purposes, 50 μm sections of the medullary pyramids were imaged separately on a Zeiss ApoTome.2 microscope. Prior to analysis, maximum intensity projections were generated for all image stacks using Fiji/ImageJ.

All axon counting was performed by three observers who were blinded to the experimental conditions, and the results were averaged for each animal. For the pyramidotomy model, a midline was drawn through the central canal of each section, and the number of GFP-positive (GFP+) axons crossing three parallel 10 μm-wide lines at 200, 400, and 600 μm lateral to the midline was counted. The number of axons was adjusted based on the total number of labeled axons passing through the medullary pyramid sections to account for differences in the tracer.

### 2.1 Workflow

The network architecture of DeNAT is illustrated in Figure 1. To preserve the fine structural details of neurites, images were not down sampled to resolutions commonly used for image classification (for example, 224 × 224 in implementations of the VGG16 architecture;(Liu and Deng, 2015)). Instead, the original resolution of 4250 × 3350 pixels was maintained throughout preprocessing. Subsequently, to facilitate an efficient training process, all images were converted to grayscale and their colors were inverted.

**Figure 1:**
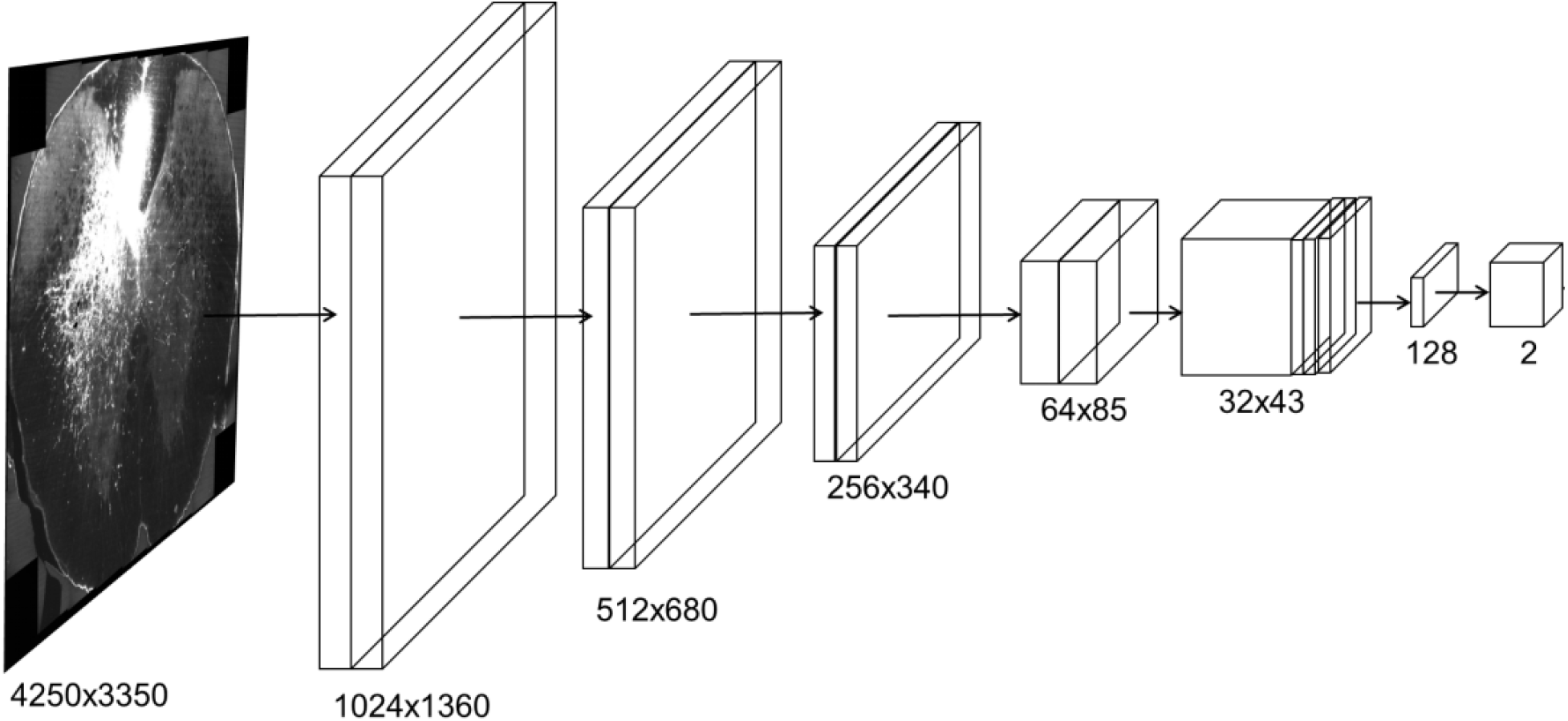
Architecture of DeNAT.

The core of the network consists of a series of paired convolutional layers. Each pair includes one strided and one non-strided convolution. The strided convolution uses a 2 ×2 stride, reducing the spatial dimensions by half at each step. The first pair of layers uses 4 filters, with the number of filters doubling at each subsequent pair up to a maximum of 64. The first convolutional layer applies a 5 ×5 kernel, while all subsequent layers use 3 ×3 kernels.

To capture information at multiple scales, side branches are generated after each main-path convolution using max-pooling. Each side branch passes through a 3 ×3 convolution with 16 filters, and the outputs are resized to a common dimension before being merged back with the main path. This design allows the model to integrate fine neurite details with broader contextual information.

Following feature aggregation, a 128-filter convolution is applied. The resulting feature maps are condensed by global average pooling into a 128-dimensional feature vector. This vector is passed to a fully connected layer with 128 nodes, followed by a 20% dropout to reduce overfitting. The final fully connected layer outputs two values, with sigmoid activation to constrain predictions between 0 and 1. All other layers are followed by ReLU activation and batch normalization.

### 2.2 Data Training

The training procedure for DeNAT is summarized in Figure 2. We used five-fold cross-validation, splitting the dataset into five equal parts. In each round, four parts (80%) were used for training and one part (20%) was reserved for testing, ensuring that the model was never evaluated on data it had already seen. To increase the effective size and diversity of the training set, we applied image augmentation. Each image was randomly flipped horizontally or vertically, and randomly shifted by up to 60 pixels along either axis, with empty regions filled with black pixels.

**Figure 2.**
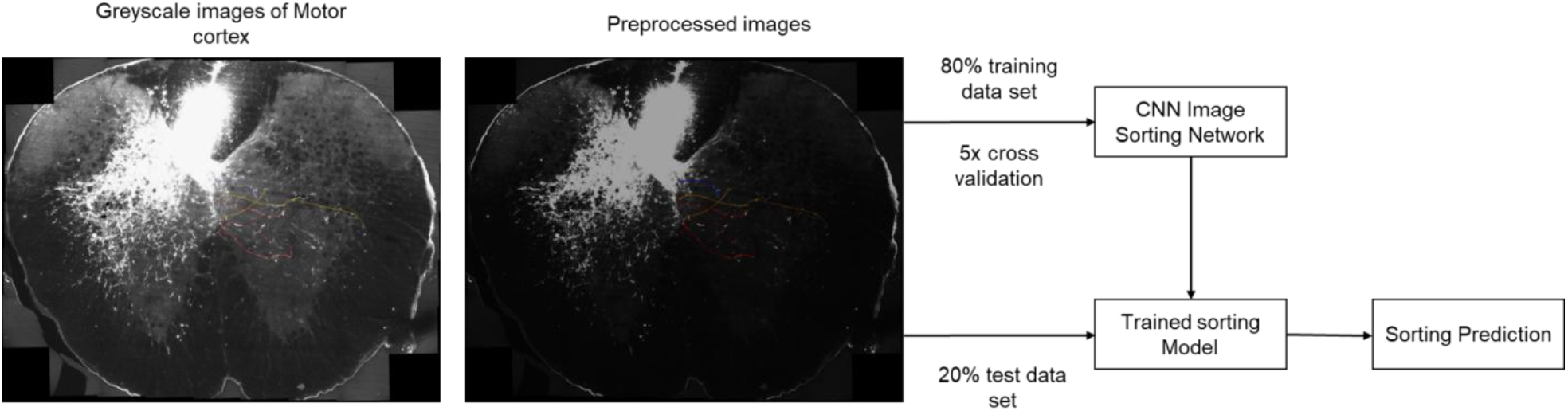
Illustrates the workflow for training and testing DeNAT.

The network was trained for 300 epochs with a batch size of 32. Binary cross-entropy loss was used, treating the two outputs as independent binary predictions. Optimization was performed using the Adam algorithm with a learning rate of 0.0001. During early experiments, we observed that the model converged faster for one output compared to the other. To balance training, we applied label smoothing, replacing strict binary labels (0 and 1) with softer targets (0.1 and 0.9). This prevented the model from overfitting to a single task while improving overall generalization.

### 2.3 Web App/Page Implementation

The DeNAT web application was designed for accessibility, allowing researchers without computational expertise to perform automated analysis with minimal setup. The interface provides simple, intuitive controls for uploading image files and options for connecting to external data sources. The front end and user interaction components were implemented using JavaScript, python, HTML and CSS, while file handling, back-end logic, and visualization modules were developed in Python. The deployed application is available at https://neuriteanalysis.netlify.app/.

The Pyramidotomy Neurite Analysis Tool interface (Figure 3) provides a streamlined workflow for analysing regenerating axons. A drag-and-drop panel allows users to upload images in PNG or JPG format, which are then processed automatically. The results are displayed as neurite counts binned by distance from the lesion site (0–200 µm, 200–400 µm, 400–600 µm, 600–800 µm, and >800 µm), enabling users to quantify sprouting with spatial precision. Output options include generating a full analysis report, exporting quantitative data in CSV format for downstream analysis, and saving labelled images for visualization and record keeping. This design ensures that both raw numerical data and processed visual outputs are available within a single, user-friendly platform.

**Figure 3:**
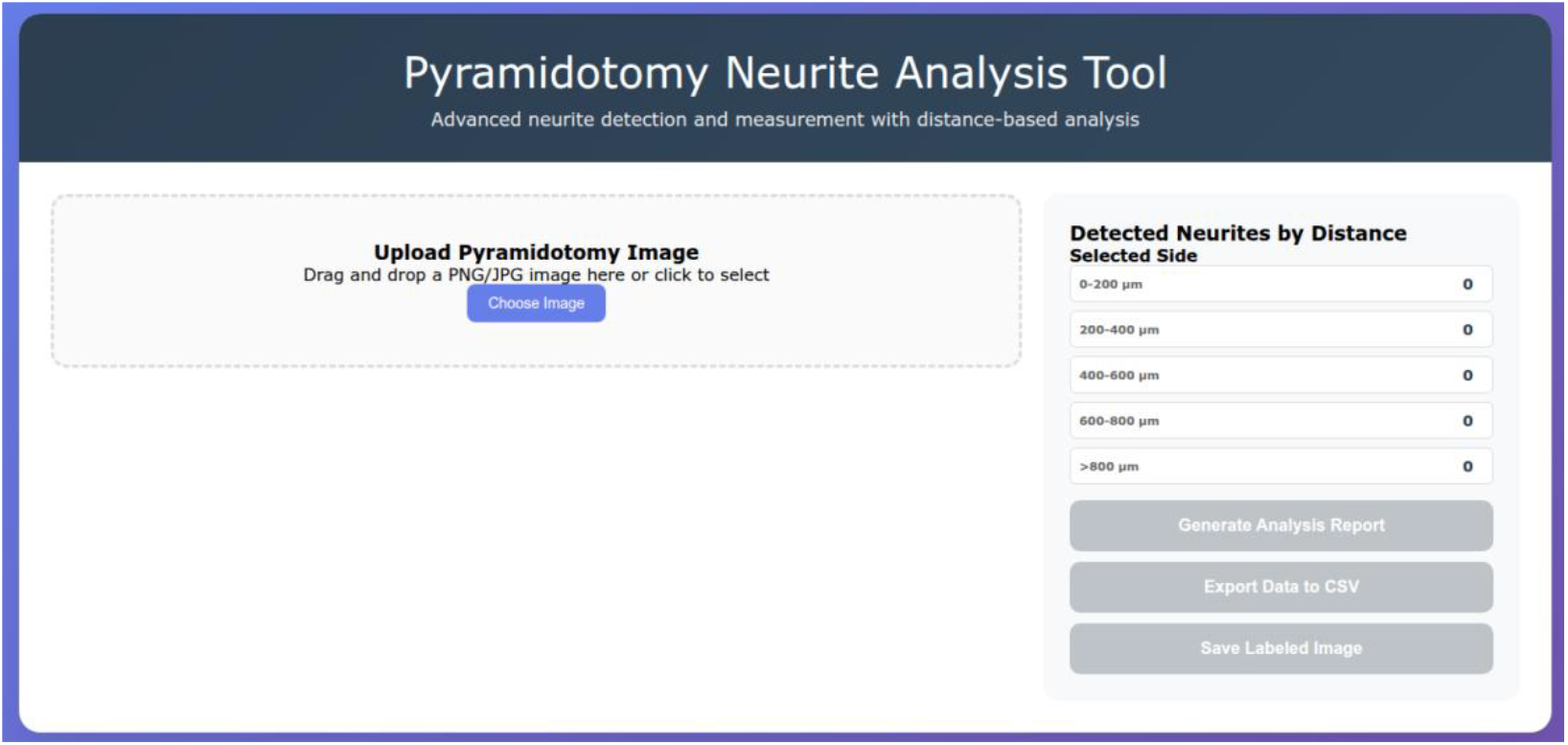
DeNAT home Page

### 2.4 Human Benchmarking Test

To evaluate DeNAT’s performance, we compared its ability to classify images against that of experienced researchers. For each test, we prepared a folder containing 20 randomly selected images from each of two groups, with filenames anonymized to prevent bias. Before sorting, researchers were given a brief presentation outlining background information, key differences between the groups, and one labeled example image from each group. They were then asked to label all images in a spreadsheet by assigning each to one of the two groups. Although the folder contained equal numbers of images per group, participants were not required to distribute their labels evenly. After completion, the true labels were revealed and researcher accuracy was scored and compared with DeNAT.

### 2.5. Statistical analyses

To compare the classification accuracy of DeNAT with that of human researchers, we first generated confusion matrices for each test set. These matrices provided counts of true positives, false positives, true negatives, and false negatives, which were used to evaluate both sensitivity and specificity. From these values, we calculated Cohen’s Kappa statistic, a measure of inter-rater agreement that accounts for agreement expected by chance (McHugh, 2012). Kappa values range from ™1 to 1, where 1 indicates perfect agreement, 0 reflects chance-level performance, and negative values indicate systematic disagreement.

For each fold of cross-validation, we computed Cohen’s Kappa separately for DeNAT and for each human participant, allowing direct comparison of performance (McHugh, 2012). Group-level averages and standard deviations were then calculated across all folds. To assess whether DeNAT significantly outperformed human raters, we evaluated its performance using standard classification metrics. Sensitivity was defined as TP /(TP + FN), precision as TP /(TP + FP), false discovery rate (FDR) as FP /(TP + FP), and the F-score as 2TP /(2TP + FP + FN). Here, TP (true positives) represent neurites that were present in the manual dataset and correctly detected by DeNAT or other tools. FP (false positives) are structures not present in the manual dataset but incorrectly identified as neurites. FN (false negatives) are neurites present in the manual dataset but not detected. Among these, the F-score was used as the primary metric for evaluating overall performance, as it integrates both sensitivity and precision into a single measure.

## 3. Results

We developed DeNAT, a novel and user-friendly tool for the quantification of neurite length. A key advantage of DeNAT over existing platforms such as ImageJ (Schneider et al., 2012), NeuronJ (W. Zhang et al., 2007), NeuriteNet (Vecchi et al., 2021), and Image-Pro Plus (Pool et al., 2008) is the integrated Region of Interest (ROI) selection feature. This enables researchers to directly define a specific area within an image and quantify neurites only within that region. By removing the need for preprocessing steps such as manual cropping, DeNAT streamlines analysis and reduces workflow inefficiency. Its most critical innovation lies in this ROI-guided approach, which addresses two major challenges in the field: the lack of biological specificity in automated quantification and the time burden associated with manual preprocessing.

As a web-based application, DeNAT also removes significant technical barriers. Unlike standalone software, it does not require local installation, dependency management, or operating system compatibility checks. Researchers can access DeNAT through a standard web browser, making it broadly accessible. In addition to automated quantification, DeNAT includes tools for manual neurite counting, image rotation, and adjustments to brightness and contrast, offering a comprehensive and flexible platform for neurite analysis.

DeNAT was developed and validated using a dataset of 231 pyramidotomy images. The dataset was partitioned into a training set (80%, 185 images) and a testing set (20%, 46 images). To generate robust reference data, three independent raters manually traced an additional validation set of 42 images. These manual annotations served as ground truth for benchmarking DeNAT’s performance against existing tools, including ImageJ with the NeuronJ plugin, Image-Pro Plus, and NeuriteNet. In this comparative analysis, DeNAT demonstrated superior performance, achieving an average accuracy of 0.77 and an F1-score of 0.86, with a False Discovery Rate (FDR) of 0.22 (Table 1). These results underscore the precision and reliability of DeNAT for automated neurite quantification in spinal cord injury models.

**Table 1:** Specificity and sensitivity of CNN vs ResNET50.

|  | TP | FP | FN | TN | Sensitivity | Precision | F-score | FDR |
| --- | --- | --- | --- | --- | --- | --- | --- | --- |
| **CNN** | **42** | **8** | **5** | **7** | **0.89** | **0.88** | **0.87** | **0.11** |
| **ResNET50** | **37** | **7** | **5** | **13** | **0.84** | **0.84** | **0.86** | **0.16** |

While we would have liked to test DeNAT against published datasets from other groups, the raw image data required for such benchmarking are rarely available. Most studies reporting pyramidotomy results present only the fiber index, defined as the ratio of axons showing cross-midline sprouting normalized to the number of labeled fibers in the medulla. As a result, we were unable to directly benchmark DeNAT using external datasets. However, if such data become available, we are eager to evaluate and validate DeNAT’s performance on independent sources.

### 3.1 CNN vs ResNet50

The performance of CNN and ResNet50 within the DeNAT framework was compared using Receiver Operating Characteristic (ROC) curves (Figure 4) and confusion-matrix–based metrics (Table 1). Both models demonstrated strong classification ability, with clear separation between sensitivity and false positive rate. The CNN model (blue curve) achieved a slightly higher Area Under the Curve (AUC) of 0.846 compared to 0.82 for ResNet50 (red curve), indicating marginally better overall discriminative performance.

**Figure 4:**
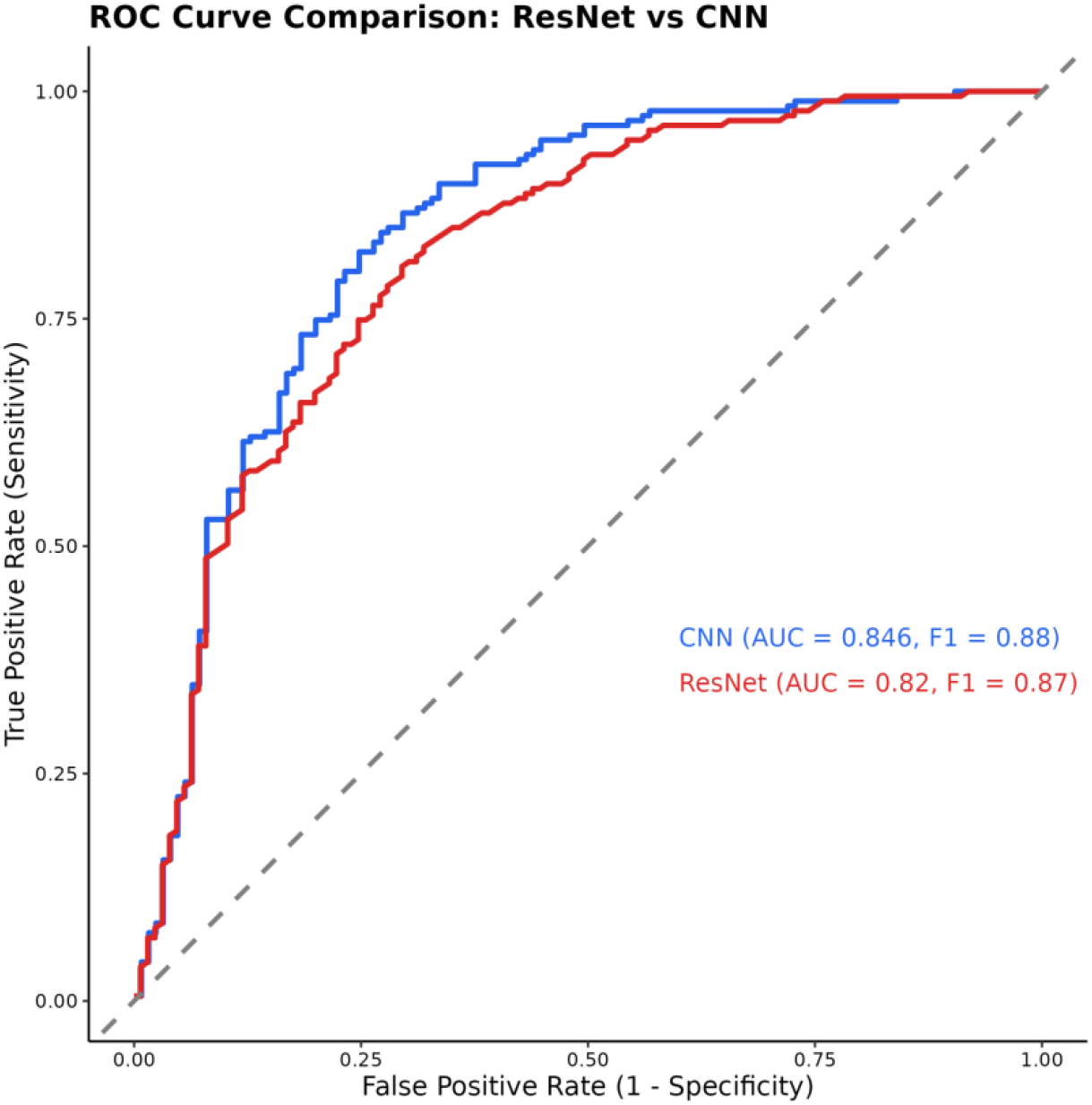
ROC curve for two different model CNN and ResNET

In terms of balanced accuracy, CNN also showed a small but consistent advantage. Its F1-score was 0.88, compared to 0.87 for ResNet50, reflecting a better balance between precision and recall. Sensitivity and precision values for CNN (0.89 and 0.88, respectively) were higher than those of ResNet50 (0.84 and 0.84). These gains translate to CNN detecting more true positives while maintaining fewer false positives. Specificity and false discovery rate (FDR) further highlight this trend. CNN achieved a specificity of 0.89 and FDR of 0.11, whereas ResNet50 had a specificity of 0.84 and a higher FDR of 0.16. Although both models performed well, CNN consistently outperformed ResNet50 across all evaluated metrics.

Taken together, these results suggest that CNN is the more reliable architecture for DeNAT, offering a modest but reproducible improvement in classification performance. Importantly, the relative simplicity of CNN compared to deeper models like ResNet50 also makes it computationally efficient, which is advantageous for deployment in user-facing applications.

### 3.2 Benchmarking DeNAT Against Established Neurite Analysis Tools

To evaluate its performance, we compared DeNAT with three widely used neurite tracing platforms: NeuronJ, Image-Pro Plus, and NeuriteNet. The benchmarking used four key metrics: precision, sensitivity, F-score, and false discovery rate (FDR) (Table 2). DeNAT achieved perfect sensitivity (1.0), consistently detecting all neurite structures present in the dataset. By contrast, NeuronJ (0.59), Image-Pro Plus (0.65), and NeuriteNet (0.70) showed substantially lower sensitivity, indicating that they missed a significant proportion of neurites. This highlights DeNAT’s strength in comprehensive detection, which is particularly advantageous in contexts where undercounting could bias results, such as regenerative growth studies or compound screening.

**Table 2:**
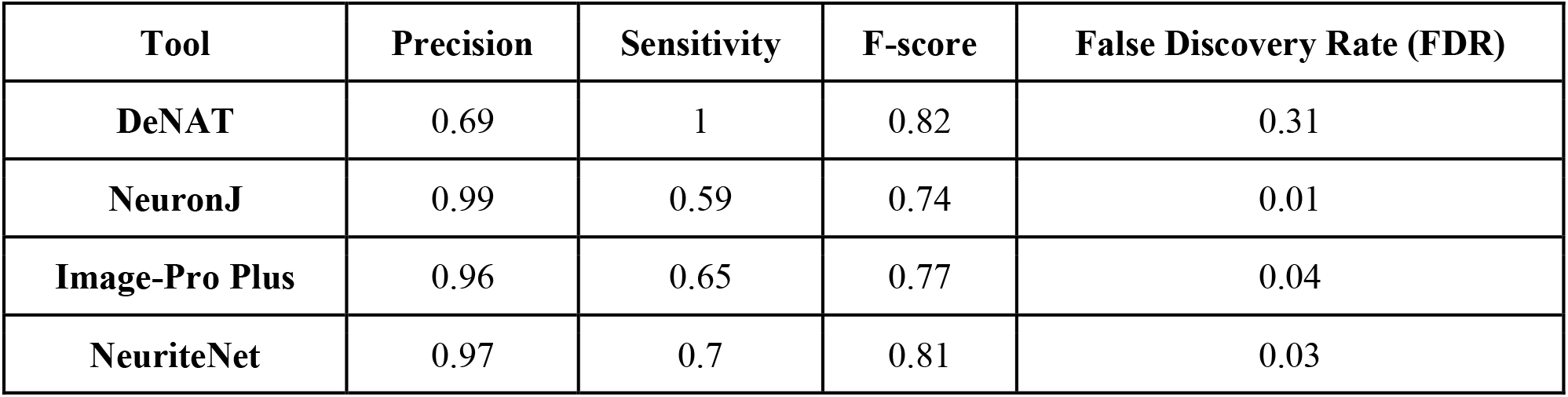
Benchmarking DeNAT Against Established Neurite Analysis Tools.

| Tool | Precision | Sensitivity | F-score | False Discovery Rate (FDR) |
| --- | --- | --- | --- | --- |
| DeNAT | 0.69 | 1 | 0.82 | 0.31 |
| NeuronJ | 0.99 | 0.59 | 0.74 | 0.01 |
| Image-Pro Plus | 0.96 | 0.65 | 0.77 | 0.04 |
| NeuriteNet | 0.97 | 0.7 | 0.81 | 0.03 |

However, this gain in sensitivity came at the cost of precision. DeNAT’s precision (0.69) was lower than NeuronJ (0.99), Image-Pro Plus (0.96), and NeuriteNet (0.97), reflecting a higher incidence of false positives. This trade-off suggests that DeNAT is more prone to over-segmentation or misclassification of background structures as neurites. Despite this, its overall accuracy, represented by the F-score (0.82), was superior to NeuronJ (0.74) and Image-Pro Plus (0.77), and closely matched NeuriteNet (0.81).Importantly, the advantage of DeNAT lies in its ability to maximize neurite detection while maintaining a competitive overall accuracy profile.

The FDR results further illustrate this balance. DeNAT had a higher FDR (0.31) than NeuronJ (0.01), Image-Pro Plus (0.04), and NeuriteNet (0.03), confirming that a larger proportion of its detections were false positives. Nevertheless, in biological applications where missing true neurites could underestimate growth, the acceptance of a higher FDR may be justified.

In summary, DeNAT maximizes neurite detection and delivers an F-score on par with the best available tools. While its precision and FDR are less favorable, its ability to avoid false negatives and capture the full extent of neurite structures makes it a robust and reliable tool for comprehensive quantification in injury and regeneration studies.

## 4 Discussion

In this study, we developed and validated the Deep Neurite Analysis Tool (DeNAT), a machine learning framework created to address a critical bottleneck in neuroscience research: the accurate and accessible quantification of neurite sprouting in complex injury models. The primary challenge with existing tools is not simply automation, but their inability to perform targeted analysis within specific anatomical regions. DeNAT overcomes this limitation with its integrated Region of Interest (ROI) selection feature, enabling researchers to quantify post-injury axonal growth while excluding irrelevant structures. Furthermore, by deploying DeNAT as a freely accessible web application, we have eliminated the high costs and computational barriers that limit the use of commercial software and advanced tracing algorithms.

Our benchmarking revealed a key trade-off that defines DeNAT’s contribution. DeNAT achieved a perfect sensitivity of 1.0, a metric where all other tested platforms underperformed. This unique ability to detect every true neurite ensures that no growth is overlooked, which is particularly advantageous in discovery-driven contexts such as high-throughput screens for neurite-promoting compounds or *in vivo* studies of subtle regenerative responses. In such settings, false negatives carry greater risk than false positives, as they could lead to the premature dismissal of promising interventions. By capturing the full landscape of neurite outgrowth, DeNAT provides a complete and more unbiased dataset for initial analysis.

We acknowledge that this exceptional sensitivity comes at the expense of lower precision (0.69) and a higher False Discovery Rate (FDR, 0.31). This reflects a greater tendency to identify background structures as neurites compared with the more conservative approaches of NeuronJ,

Image-Pro Plus, and NeuriteNet. Nonetheless, we argue that this trade-off is acceptable in many biological applications. False positives can often be corrected through manual review, whereas false negatives cannot be recovered once missed. DeNAT’s robust F1-score of 0.82 underscores that it achieves competitive overall performance by balancing sensitivity with reasonable accuracy.

The technical foundation of DeNAT—a custom CNN architecture trained on high-resolution images without downsampling—likely underpins its ability to capture faint or fine neurite extensions that other methods overlook. Future iterations of DeNAT will prioritize improving precision without compromising sensitivity. This may be achieved by expanding the training dataset to include a wider range of neuronal types, injury paradigms, and imaging conditions, or by integrating post-processing filters to reduce artifacts. While DeNAT was validated primarily on the pyramidotomy model, we plan to extend its application to the thoracic crush model, which will broaden its relevance to additional spinal cord injury paradigms.

A current limitation of this work is that benchmarking was conducted only on our own datasets. Ideally, we would have compared DeNAT against raw data from other published pyramidotomy studies. However, such datasets are rarely made publicly available; most reports present only derived values such as the fibre index (the ratio of axons crossing the midline, normalised to labelled fibers in the medulla). Without access to underlying images, external benchmarking is not possible. Should such datasets become available, we are eager to validate DeNAT across independent sources.In conclusion, DeNAT provides a unique combination of accessibility, flexibility, and comprehensive detection. The design prioritises sensitivity and reproducibility, offering researchers a powerful new tool for quantifying neurite sprouting in challenging injury models. With further refinement and expansion to additional paradigms such as thoracic crush, DeNAT has the potential to become a broadly applicable platform for axon regeneration research.

## Author Contributions

Manojkumar Kumaran was responsible for conceptualization, investigation, methodology, data curation, software development, formal analysis, data interpretation, and writing of the original draft. Athul Narayan P.S., Yogesh Sahu, Soupayan Banerjee contributed to data curation, data interpretation, imaging and review. Anisha S. Menon contributed to conceptualization, data interpretation, formal analysis, imaging, and review and editing. Shringika Soni contributed to imaging and data interpretation. Ishwariya Venkatesh contributed to conceptualization, investigation, methodology, and writing, review and editing.

### Data availability statement

*https://github.com/mano2991/Neurite-Analysis-Tool*

*https://neuriteanalysis.netlify.app/*

### Funding

This research was supported by funding from Council of Scientific and Industrial Research (CSIR - HCP53201), Department of Biotechnology (DBT - BT/PR51467/MED/122/358/2024), and Science and Engineering Research Board (SER - SRG00116).

## Acknowledgments

We thank CSIR-CCMB for providing research facilities and support staff. We thank Mr. B. Suman and Dr. Nitla Venkata Mahesh (Microscopy) for their technical assistance and Dr. Prakash Chermakani for reviewing and for writing suggestions.

